# Endogenous Opioid System Modulates Neural Activation During Emotion Regulation

**DOI:** 10.64898/2026.07.30.739761

**Authors:** Vesa Putkinen, Ananya Ravindran, Kerttu Seppälä, Harri Harju, Anita Song, Lauri Nummenmaa

## Abstract

Emotion regulation is critical for adaptive functioning and mental health. Functional neuroimaging studies have identified distributed frontal, parietal, and limbic networks involved in emotion regulation, particularly during cognitive reappraisal. However, the neurochemical mechanisms underlying individual differences in emotion regulation remain poorly understood. The endogenous opioid system is a promising candidate because μ-opioid receptor systems acts as a stress buffer and modulates emotional responses to both rewarding and aversive stimuli

We used positron emission tomography (PET) and functional magnetic resonance imaging (fMRI) to examine whether baseline μ-opioid receptor (MOR) availability is associated with haemodynamic responses during emotion regulation through reappraisal. Participants underwent [^11^C]carfentanil PET imaging to quantify MOR availability and subsequently completed an fMRI task involving passive viewing of emotional images and cognitive reappraisal of their emotions during image viewing

As expected, fMRI revealed engagement of the dorsolateral prefrontal cortex and inferior parietal lobule during cognitive reappraisal. Baseline MOR availability predicted blood oxygenation level-dependent (BOLD) responses in frontal, temporal, and parietal regions: individuals with higher MOR availability showed a larger difference between regulation and viewing trials in the temporal pole, angular gyrus, anterior insula, thalamus, and inferior frontal gyrus. These associations were driven primarily by stronger responses during passive viewing, although some regions also showed evidence of reduced responses during active reappraisal.

These findings suggest that higher endogenous opioid tone is associated with stronger recruitment of distributed frontal, temporal, and parietal networks during emotional stimulus processing as well as attenuation of activity in some of these regions during cognitive reappraisal. Thus, while higher MOR availability is associated with enhanced neural responsiveness to emotional stimuli, these responses can nevertheless be effectively modulated through deliberate emotion regulation.

## Introduction

Emotions guide adaptive behaviour by signaling the motivational significance of events and promoting approach toward rewards and avoidance of threats. At the same time, transient negative emotions can undermine motivation, impair decision-making and sustained negative affect may compromise immune function, predispose to cardiovascular diseases and increase all-cause mortality (Suinn 2001). Even positive emotions can have maladaptive consequences, such as increased risk taking or judgment biases, and may be inappropriate in certain contexts (e.g., funerals). Accordingly, emotion regulation is central to maintaining well-being, pursuing long-term goals, and navigating social relationships and difficulties in emotion regulation or reliance on maladaptive regulation strategies (e.g., alcohol use) are linked to poorer academic and occupational performance and compromised psychological and physical health (Lincoln et al. 2022).

Emotion regulation can be achieved through a variety of strategies targeting different stages of the emotion-generating process, ranging from selecting emotion-eliciting situations or stimuli to reappraising their meaning or suppressing the outward expression of emotion (Gross 1998). Cognitive reappraisal refers to altering the emotional impact of a stimulus or situation, for instance by reinterpreting it (e.g., “The shocking scene is fake”) or distancing oneself from it (e.g., “This does not concern me”). The neural basis of cognitive reappraisal has been extensively studied with neuroimaging (Urry et al. 2006; Beauregard et al. 2001; Ochsner et al. 2002) due to its theoretical significance, clinical relevance, and the ease with which it can be operationalized in experimental research. This research has consistently implicated the dorsolateral prefrontal cortex (DLPFC), ventrolateral prefrontal cortex/inferior frontal gyrus (VLPFC/IFG), and inferior parietal lobule in the cognitive reappraisal of both positively and negatively valenced stimuli (for meta-analyses, see (Buhle et al. 2014; Morawetz et al. 2017). Thus, cognitive reappraisal engages a fronto-parietal network that is broadly involved in domain-general cognitive control processes, including working memory, attentional control, and response inhibition.

While the neural circuits supporting emotion regulation have been extensively studied, the underlying neuromolecular mechanisms remain poorly understood. The μ-opioid receptor (MOR) system may contribute to emotion regulation given its well-established involvement in pleasure, pain, and emotion more broadly (Leknes and Tracey 2008; Nummenmaa and Tuominen 2018). Positron emission tomography (PET) studies have shown that rewarding stimuli such as food, social touch, sexual activity, and music modulate MOR activity (Jern et al. 2023; Putkinen et al. 2025), while pharmacological down- and upregulation of the MOR system alters reward, threat, and stress responses (Meier et al. 2021). Individual differences in baseline MOR availability are associated with haemodynamic responses to both pleasant and unpleasant stimuli, suggesting that the MOR system broadly modulates emotional processing (Putkinen et al. 2025; Seppälä et al. 2025; Sun et al. 2022; Karjalainen et al. 2019; Nummenmaa et al. 2018). In addition, MORs have been implicated in cognitive processes relevant for emotion regulation such as management of stress and anxiety (Ribeiro et al. 2005; Nummenmaa et al. 2020), potentially influencing the allocation of regulatory resources through their effects on motivation and affective arousal rather than through direct modulation of executive control (van Steenbergen et al. 2019). Therefore, the MOR system represents a promising neuromolecular mechanism contributing to emotion regulation.

### The current study

Here we investigated the role of the MOR system using fusion imaging with PET and fMRI. Haemodynamic responses during emotion regulation were predicted with baseline MOR availability measured with PET using the MOR-selective radioligand [^11^C]carfentanil. During the fMRI experiment, participants viewed positive and negative images and were instructed either to regulate their emotional responses via reappraisal or to view the images and experience the evoked emotions. The fMRI results revealed that emotion regulation engaged the canonical emotion regulation regions, DLPFC and inferior parietal lobule during regulation compared to view trials. Baseline MOR availability was associated with the emotion regulation dependent haemodynamic responses in frontal, temporal and parietal regions. This effect was driven primarily by enhanced responses during view trials, while some regions also showed attenuated responses during regulation in individuals with higher MOR availability. Together, these findings suggest that individuals with higher MOR availability exhibit stronger recruitment of distributed heteromodal networks during emotional stimulus processing, yet are also able to attenuate activity within these regions via cognitive reappraisal.

## 2. Methods

### Participants

Thirty-two healthy adult women (mean age = 24 years, range = 19–42 years) participated in the study. All were right-handed, had normal or corrected-to-normal vision, and reported no history of neurological or psychiatric disorders. Only women were included to maximize statistical power, as the spatial distribution of MORs exhibits sex-related variability (Kantonen et al. 2020). Additionally, women tend to report stronger emotional responses, and a female-only sample was therefore expected to maximize task-related effects in this complex PET study (Lench et al. 2011). Fifteen of these participants underwent both PET and fMRI while the remaining participated only in the fMRI session. Eligibility for PET/MR imaging was assessed by the study physician, and psychiatric disorders were evaluated by a psychologist using the MINI 6.0 interview. Written informed consent was obtained prior to participation. The study was approved by the ethics board of the Hospital District of Southwest Finland and conducted in accordance with the Declaration of Helsinki.

### Stimuli

The stimuli consisted of 35 aversive and 25 pleasant images. All pleasant and 25 of the unpleasant images were chosen from the Nencki Affective Picture System (NAPS; (Marchewka et al. 2014)). The pleasant images depicted, for example, affiliative social interactions and animals, whereas the aversive images included scenes of injury and threat. Additionally, 10 aversive snake images were included to accommodate a separate study protocol on fear of snakes (Seppälä et al. 2025).

### Task Design and Procedure

Before the experiment, the procedure was explained to the participants. For the emotion regulation trials, the participants were instructed to regulate their emotional responses using cognitive reappraisal. This was described as trying to view the images as less personally relevant, reinterpreting their meaning (e.g., by assuming that the depicted situation is temporary or less severe than it appears), or considering that the scenes were staged or not real. For the view trials, they were instructed to view the images and allow their emotions to arise naturally without attempting to regulate them. Participants completed a short practice session prior to scanning to ensure they understood the task.

Participants performed an emotion regulation task (**Figure 1**) during the fMRI scan. Each trial began with a fixation cross (0.5 s) followed by a cue (1 s) indicating the current instruction. A green circle cued participants to view the pictures naturally without attempting to modify their emotional response, while a red cross instructed them to actively regulate their emotions (“Regulate” condition) as described above.

**Figure 1.**
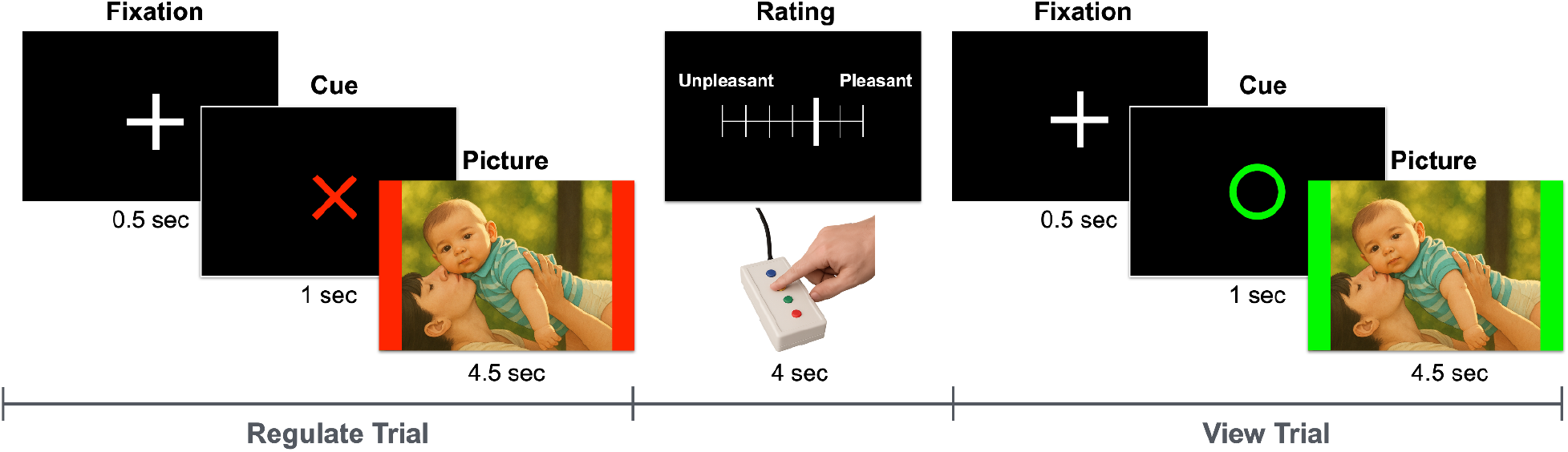
Experimental design. Each trial began with a fixation cross, followed by a cue indicating whether participants should regulate their emotional response to the subsequent picture (red “X”) or simply view it and freely feel the emotions elicited by it (green “O”). To reinforce the instruction, the picture was framed with bars matching the cue color. After each picture, participants rated their emotional response on a 7-point scale ranging from unpleasant to pleasant. On both trial types, 50% of the pictures depicted positive and 50% negative emotional content. The example stimulus is AI-generated and is provided solely for illustrative purposes to avoid including identifiable photographs of individuals in accordance with bioRxiv’s guidelines.

An emotional image (positive or negative) was then presented for 4.5 s, bordered by a color corresponding to the instruction (green = view, red = regulate). Participants were asked to remain still, attend to the image, and follow the cue by either experiencing or regulating their emotions.

After each image, participants rated their subjective affective responses to the picture on a continuous valence scale ranging from *Unpleasant* (0) to *Pleasant* (10). Ratings were made using an MRI-compatible button box by moving a cursor along the scale. Each trial lasted 6 s, and the conditions were pseudorandomized to prevent more than two consecutive trials of the same type.

### MRI Data Acquisition and Preprocessing

MRI data were acquired using a 3 T GE SIGNA Premier scanner equipped with a 48-channel head coil. High-resolution anatomical images for EPI and PET data normalisation were obtained using a T1-weighted MPRAGE sequence (1 mm^3^ isotropic resolution). Functional images were collected over 24 minutes using a T2*-weighted echo-planar imaging (EPI) sequence sensitive to BOLD contrast (TR = 2600 ms, TE = 30 ms, 45 axial slices, 3 mm thickness). A total of 470 volumes were acquired per fMRI run.

Preprocessing of the fMRI data was conducted using FMRIPREP (Esteban et al., 2019). Structural images were corrected for intensity non-uniformity and skull-stripped. Cortical surface reconstruction was performed with FreeSurfer, and spatial normalization to MNI152 standard space was achieved via nonlinear registration. Brain tissue segmentation into gray matter, white matter, and cerebrospinal fluid (CSF) was performed using FAST (FSL). The preprocessed functional data were coregistered to the corresponding T1-weighted image, resampled to MNI space, and smoothed with an isotropic Gaussian kernel of 6 mm FWHM. To minimize motion-related noise, ICA-AROMA (Pruim et al., 2015) was applied.

### PET data acquisition and processing

MOR availability was quantified using the radioligand [^11^C]carfentanil which has high affinity for mu-opioid receptors and high test-retest reliability (Hirvonen et al. 2009). The radioligand was synthesized as described previously (Kantonen et al. 2022). The radiochemical purity of the produced [^11^C]carfentanil batches was 98.1 ± 0.4% (mean ± SD). The injected radioactivity was 252 ± 10 MBq, with a molar radioactivity of 354 ± 240 MBq/nmol at the time of injection, corresponding to an injected mass of 0.52 ± 0.43 μg.

PET imaging was performed using a Discovery 690 PET/CT scanner (GE Healthcare, USA). The tracer was administered as a single intravenous bolus via a catheter placed in the antecubital vein, and radioactivity was measured for 51 minutes following injection. The participant’s head was stabilized with straps to minimize movement during scanning. Prior to PET imaging, a computed tomography scan was acquired for attenuation correction.

Preprocessing of PET data was carried out using the in-house Magia toolbox (https://github.com/tkkarjal/magia) within MATLAB (The MathWorks, Inc., Natick, MA, USA) (Karjalainen et al. 2020). PET images were first corrected for motion and coregistered to T1-weighted MR images, which were subsequently processed with FreeSurfer to derive anatomical parcellations. [11C]carfentanil binding potential (BP_ND_), relative to non-displaceable uptake, was estimated voxel-wise using the simplified reference tissue model (SRTM) with the occipital cortex as the reference region. The resulting parametric images were spatially normalized to MNI152 space based on the warping of the T1 images and smoothed using a Gaussian kernel with 6 mm FWHM.

### Statistical Analysis

First-level analyses for fMRI data were performed in SPM12 using the General Linear Model (GLM). Each participant’s time series was modeled with four regressors representing the four experimental conditions (View positive, Regulate positive, View negative, Regulate negative Regulation). Each regressor was modeled as a boxcar function corresponding to the 4.5 s stimulus presentation period and convolved with the canonical hemodynamic response function (HRF).

First-level contrasts of interest included View > Regulate, Regulate > View, View positive > Regulate positive, and View negative > Regulate negative. The resulting contrast images from each participant were entered into random-effects group-level analyses, implemented as one-sample *t*-tests in SPM12.Group-level inference was performed using a voxel-wise threshold of p < 0.001 (uncorrected), followed by cluster-level FWE correction at p < 0.05.

The association between baseline MOR availability and BOLD responses during the emotion regulation task was quantified through whole-brain correlational analysis. Because [11C]carfentanil binding has high regional autocorrelation (Tuominen et al. 2014), we first performed a principal component analysis (PCA) on the mean carfentanil binding in regions defined by the AAL atlas (excluding the cerebellum) and subsequently correlated the scores for the first principal component (explaining 70% of the variance in the region-wise MOR availability) with the voxel-wise beta-values for the BOLD contrasts of interest (see **Figure S1** for an illustration of regional loadings for the first PC). As this analysis was restricted to the 15 participants who underwent both fMRI and PET, and therefore had substantially lower statistical power, the results are reported with a more lenient voxel-wise threshold of *p* < 0.01 (uncorrected), with cluster-level FWE correction at *p* < 0.05.

As a follow-up to the whole-brain analysis, clusters showing significant associations between baseline MOR availability and the Regulate > View contrast in fMRI were used as regions of interest. Mean BOLD responses for the Regulate and View conditions were then extracted separately from these regions and correlated with baseline MOR availability. This analysis was performed to characterize the contributions of each condition to the whole-brain Regulate > View contrast rather than to provide independent statistical inference.

## 3. Results

### Subjective pleasantness ratings

A 2 × 2 repeated-measures ANOVA with Stimulus Type (Positive vs. Negative) and Condition (Regulate vs. View) revealed a robust main effect of Stimulus Type, F(1,32) = 698.90, p < .001, η^2^ = .89, confirming that positive pictures were rated as more pleasant than negative pictures. A main effect of Condition was observed, F(1,32) = 4.70, p = .038, η^2^ = .018, but this effect was qualified by a strong Stimulus Type × Condition interaction, F(1,32) = 170.71, p < .001, η^2^ = .56. Follow-up inspection of the interaction indicated that, relative to Regulation trials, ratings in the View condition were more positive for the positive stimuli and more negative for negative stimuli, indicating that the effect of Condition depended strongly on stimulus valence (**Figure 2**).

**Figure 2.**
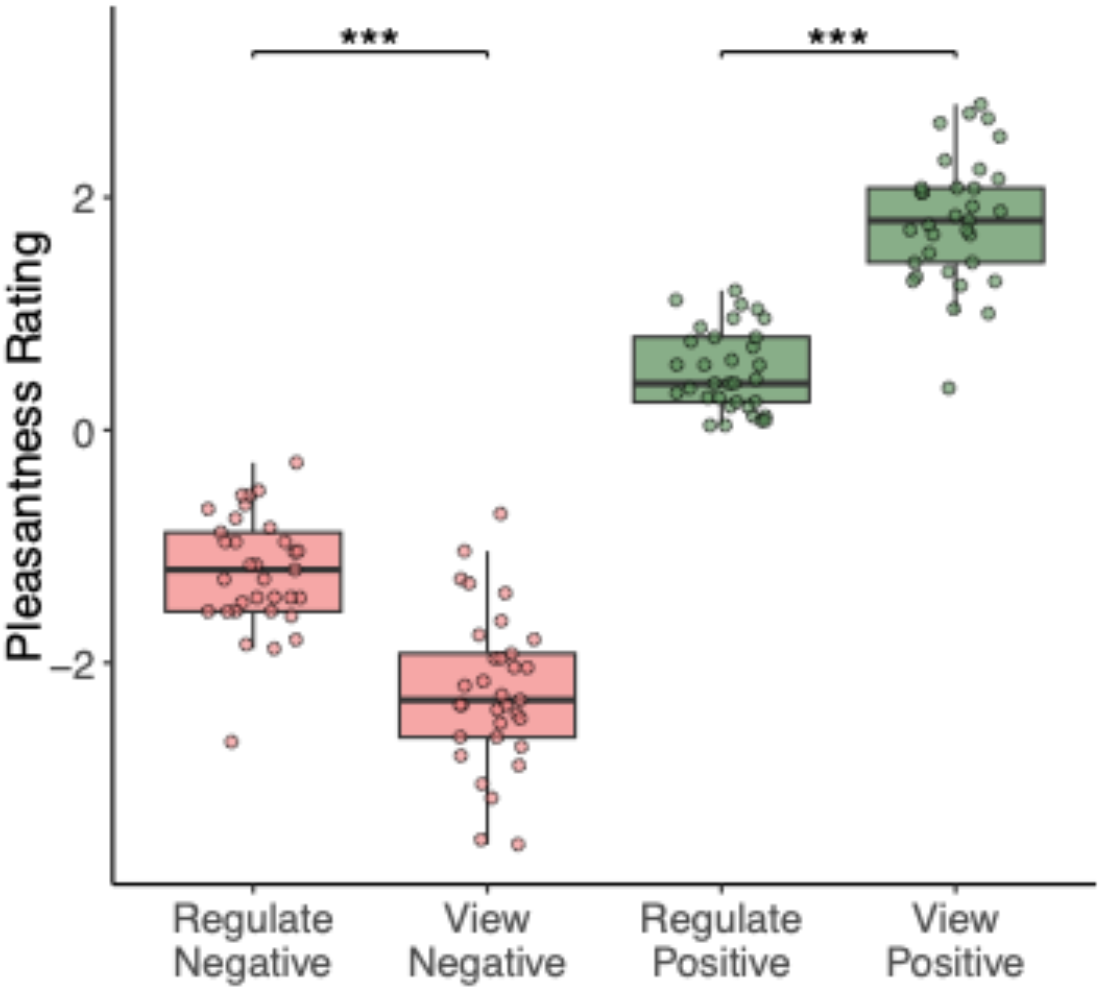
Pleasantness ratings collected during fMRI scanning for positive and negative images in the View and Regule conditions. Ratings were made on a scale from −3 (very unpleasant) to +3 (very pleasant). *** = p < .001

### fMRI results

### Main effect of emotion regulation

The Regulate > View contrast revealed stronger activation in the bilateral inferior parietal lobule (IPL) and the right dorsolateral prefrontal cortex (DLPFC) (**Figure 3**). The opposite contrast showed greater activation in the postcentral gyrus, supplementary motor area (SMA), insula, thalamus, and lateral occipital and and superior parietal cortex. For valence specific effects, see supplementary material (**Figure S2**)

**Figure 3.**
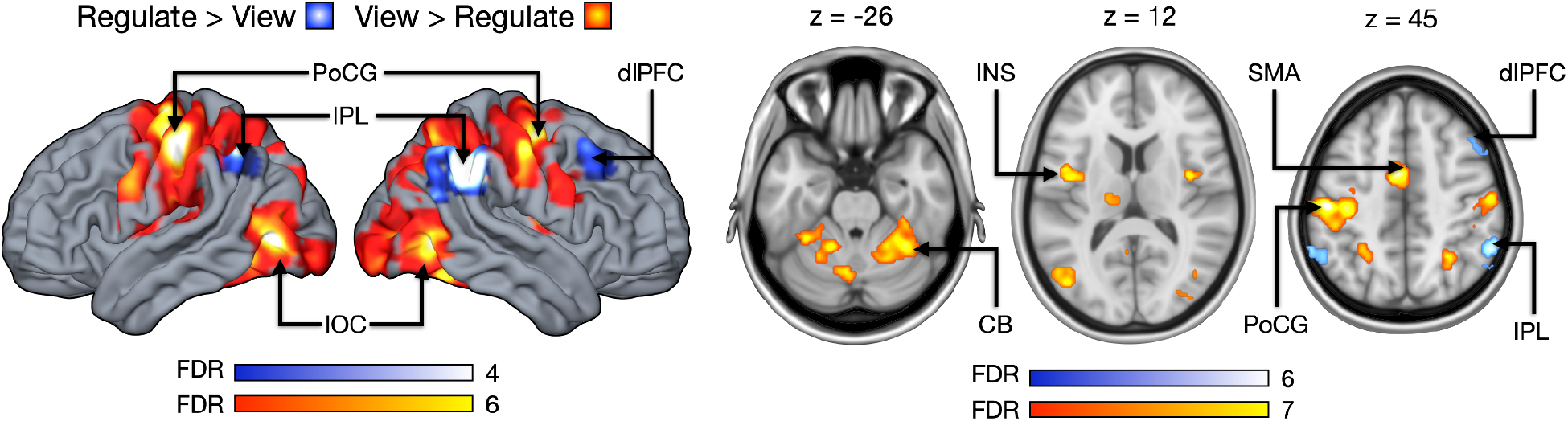
Regions showing increased BOLD responses for the Regulate > View and View > Regulate contrasts. Statistical maps are thresholded at a voxel-wise p < 0.001 (uncorrected) and cluster-level FWE-corrected at p < 0.05. The color bar indicates t-value. CB = cerebellum, dfPFC = dorsolateral prefrontal cortex, INS = insula, IPL = inferior parietal lobule, lOC = lateral occipital cortex, PoCG = postcentral gyrus, SMA = supplementary motor area.

### fMRI-PET fusion analysis

Finally, we assessed whether baseline MOR availability was related to whole-brain BOLD responses during emotion regulation. Higher MOR availability was significantly associated with stronger hemodynamic responses for the Regulate > View contrast in inferior frontal cortex (IFG), angular gyrus, temporal pole and the thalamus (**Figure 4**). No significant negative associations between MOR tone and Regulate> View BOLD contrast were observed.

**Figure 4.**
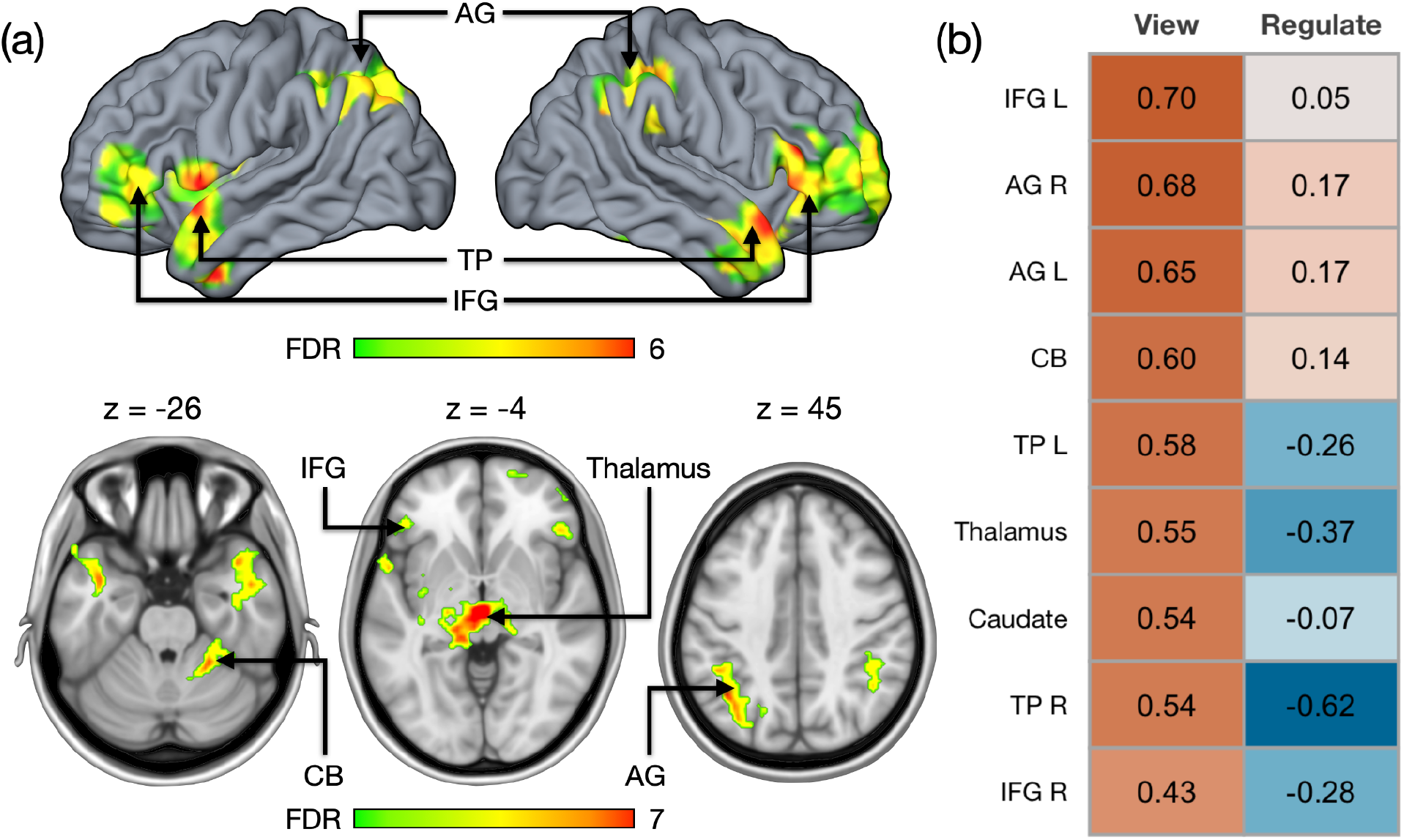
(a) Regions where baseline [^11^C]carfentanil binding was positively associated with BOLD activation for the Regulate > View contrast. Statistical maps are thresholded at a voxel-wise p < 0.01 (uncorrected) and cluster-level FWE-corrected at p < 0.05. The color bar indicates t-value. (b) Correlations between baseline BP_ND_ and mean beta values in selected ROIs for the View and Regulate trials. AG = angular gyrus, CB = cerebellum, IFG = inferior frontal gyrus, TP = temporal pole

## Discussion

Our main finding was that MOR availability predicted hemodynamic responses in frontal, temporal, and parietal regions during an emotion regulation task. Specifically, individuals with higher baseline MOR availability showed larger BOLD responses when regulating their emotions versus viewing the affective images passively. Follow-up analyses indicated that MOR availability was associated with both enhanced responses during passive viewing of emotional images (across the regions) and attenuated responses during emotion regulation in IFG, temporal pole and thalamus. Together, these findings suggest that baseline MOR availability is associated with bidirectional modulation of distributed cortical systems engaged during emotional processing and cognitive reappraisal.

### Baseline MOR availability predicts hemodynamic responses during natural viewing and reappraisal

Our findings extend previous PET-fMRI studies showing that baseline MOR availability predicts hemodynamic responses to emotional stimuli (Putkinen et al. 2025; Seppälä et al. 2025; Nummenmaa and Tuominen 2018). Although the larger Regulate > View differences in individuals with higher MOR availability were driven primarily by stronger responses during passive viewing, some regions, particularly the temporal pole, also exhibited negative associations between MOR availability and BOLD responses during regulation, further accentuating this difference (**Figure 4b and S3**). This finding indicates that individuals with higher MOR availability exhibit heightened neural responsiveness to emotional stimuli yet also show attenuation of activity within portions of the same network during cognitive reappraisal.

MOR-BOLD associations were observed in thalamus and caudate nucleus, but also in parietal, temporal and frontal regions that fall outside the canonical emotion circuits. Parietal activity was found in the angular gyrus, which has been described as a multimodal association hub with several functions relevant to the processing of emotional visual stimuli, including attention reorienting, visual search, social cognition, self-processing, semantic processing, and episodic/autobiographical memory retrieval (Seghier 2023). The temporal effects were concentrated on the temporal pole which has also been implicated in emotional processing, theory of mind, autobiographical memory, semantic processing, and integration of socially relevant multimodal information (Herlin et al. 2021). An association between MOR availability and BOLD responses was also observed in a left-lateralized cluster spanning portions of the anterior insula, frontal operculum, and inferior frontal gyrus. The anterior insula, in particular, has been implicated in emotional processing, arousal, and interoception (Gu et al. 2013; Craig 2009). Together, these findings suggest that individuals with higher MOR availability engage distributed heteromodal cortical systems more strongly during affective processing, whereas MOR-dependent variation was less evident within the core limbic circuitry.

### Neural circuits supporting emotion regulation

The fMRI results revealed that reappraisal engages the dlPFC and inferior parietal lobule. These regions are involved in domain-general cognitive control, attention and working memory processes that support deliberate emotion regulation (Toh et al. 2024; Buhle et al. 2014). Although prior studies have frequently reported IFG engagement during reappraisal tasks (Morawetz et al. 2017), we did not observe IFG recruitment in the Regulate > View contrast. The IFG has also been implicated more broadly in semantic processing and evaluative appraisal during emotional tasks (Morawetz et al. 2017). Thus, participants may have spontaneously engaged such processes during unconstrained viewing of emotional stimuli reducing the differential IFG response between regulate and view conditions. This interpretation is further supported by the observation that individuals with higher MOR availability showed stronger IFG responses during view trials, suggesting greater spontaneous engagement of appraisal- or semantic-related processing even in the absence of explicit regulation instructions. We did, however, observe IFG activation for the Regulate > View contrast when the analysis was restricted to positive images only (**Figure S2**). Although speculative, it is possible that the more aversive negative images elicited spontaneous recruitment of regulatory processes during viewing, thereby reducing the additional IFG engagement required during explicit reappraisal and consequently attenuating the overall Regulate > View contrast when positive and negative images were analyzed together.

The View > Regulate contrast revealed increased activation in lateral occipital regions, consistent with enhanced visual processing of emotionally salient stimuli. Enhanced responses were also observed in somatosensory cortex, insula, and SMA, suggesting increased bodily/interoceptive processing and autonomic engagement during unconstrained emotional viewing. Together, these findings indicate that the emotional stimuli primarily modulated sensory and embodied aspects of affective processing rather than strongly engaging the subcortical reward or salience regions, as we did not observe robust ventral striatal or amygdala responses in the View > Regulate contrast despite evidence that reappraisal frequently modulates amygdala activity (Buhle et al. 2014). This suggests that the present task primarily engaged perceptual and bodily aspects of emotional processing rather than strong differential limbic responding between conditions.

### Limitations

We recruited only women to minimize sex-dependent variation in MOR availability and to maximize statistical power (Kantonen et al. 2020). The findings may thus not generalize to men. Although the task included both positive and negative emotional pictures, the study was underpowered to examine valence-specific effects of baseline MOR availability on BOLD responses because the PET-fMRI analyses were restricted to the 15 participants who underwent both PET and fMRI, requiring subdivision of trials into four conditions. While cognitive reappraisal is thought to rely on broadly overlapping neural mechanisms across emotional valence, possible valence-dependent differences in MOR-related effects remain to be investigated. [^11^C]carfentanil BP_*ND*_ reflects both receptor density and endogenous opioid occupancy, and the present design cannot disentangle these explanations. Finally, the study was run using cross-sectional design and it cannot reveal acute effects of emotion regulation on opioidergic neurotransmission; this effects needs to be established in future studies using repeated PET imaging with the challenge paradigm (Manninen et al. 2017; Seppälä et al. 2025; Putkinen et al. 2025)

## Conclusions

Baseline MOR availability is positively associated with neural responses in temporal, frontal, and parietal regions during an emotion regulation task. These effects reflect primarily heightened responses during natural viewing of emotional stimuli, while responses in temporal pole, IFG and thalamus are reduced during emotion regulation in individuals with higher MOR availability. Together, these findings suggest that higher MOR availability is associated with greater neural sensitivity to emotional stimuli and stronger downregulation of responses during cognitive reappraisal. Taken together, these results align with the role of the MOR system in stress buffering and affect regulation (Seppälä et al. 2025; Nummenmaa et al. 2020).

## Acknowledgements

We sincerely thank Eleni Rebelos for contributing to the PET data collection. The work was partly supported by the Academy of Finland (#350416) to VP, Jane and Aatos Erkko Foundation, and European Research Council Advanced Grant (#101141656) to LN.

## Supplementary Material

**Supplementary Figure S1.**
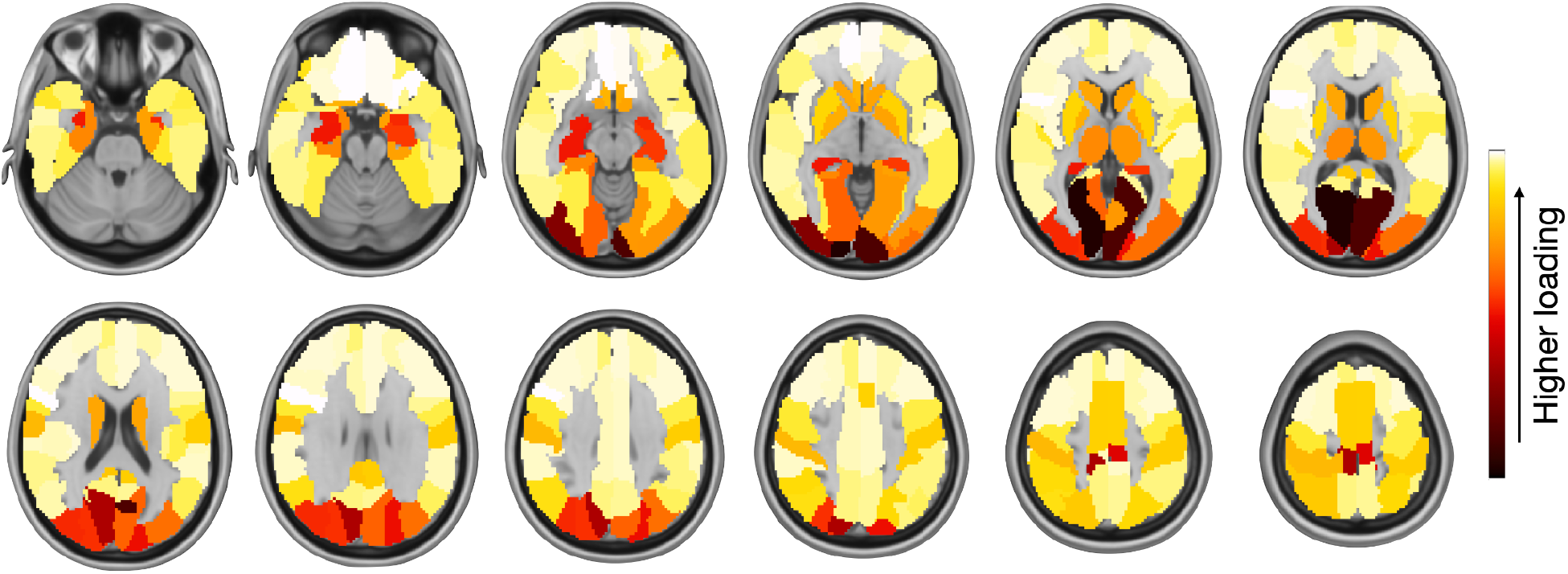
Regional loadings for the first BP_*ND*_ principal component.

**Supplementary Figure S2.**
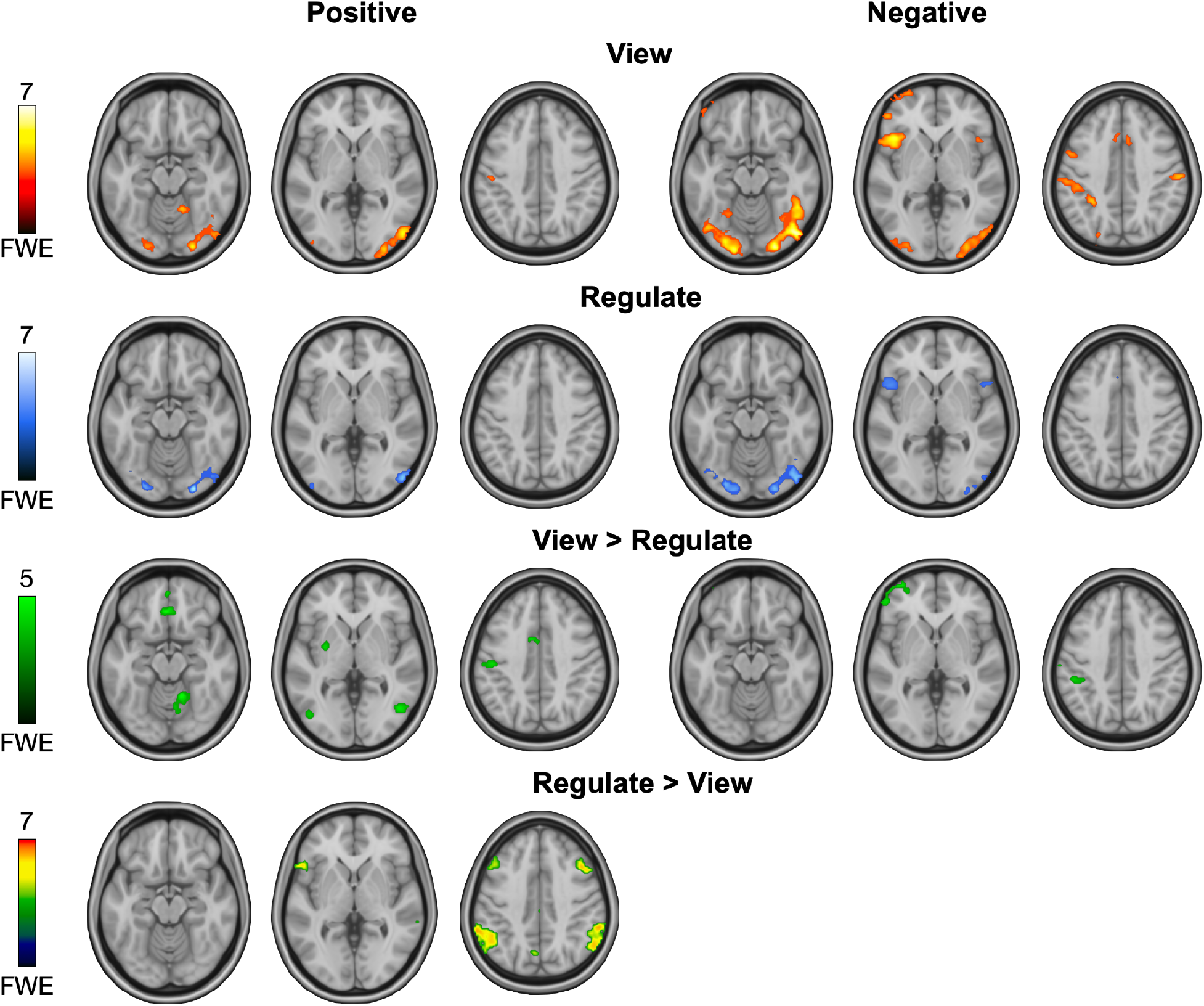
BOLD responses for the Regulate, View, View > Regulate, and Regulate > View contrasts shown separately for positive and negative images. The Regulate > View contrast for negative images yielded no significant clusters and is therefore not displayed. Statistical maps are thresholded at a voxel-wise p < 0.05 (uncorrected) with cluster-level FWE correction at p < 0.05.

**Supplementary Figure S3.**
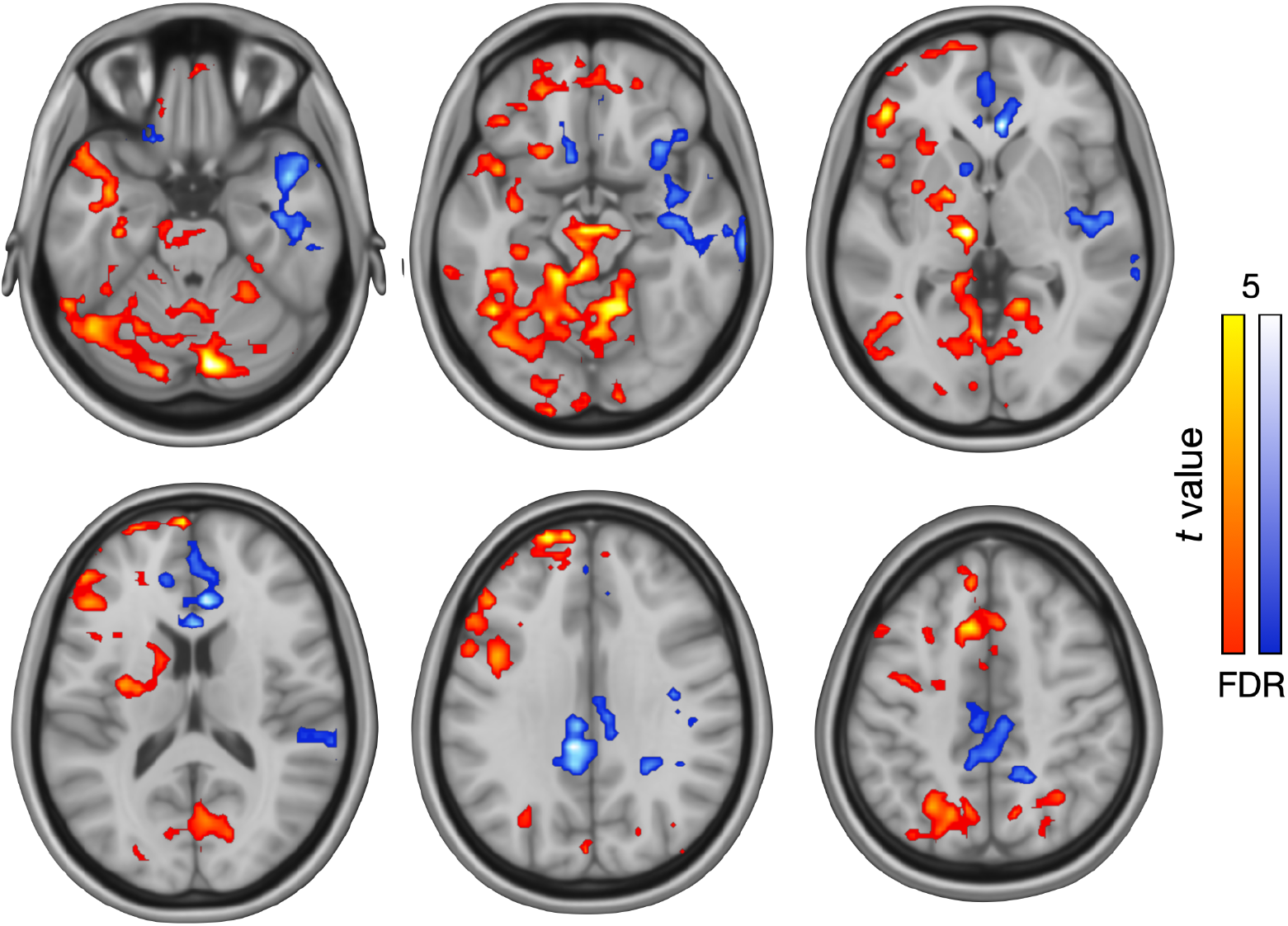
Whole-brain voxel-wise associations between the first principal component (PC1) of baseline MOR availability and BOLD responses during the View and Regulate trials.

